# Engineered cartilage from human chondrocytes with homozygous knockout of cell cycle inhibitor p21

**DOI:** 10.1101/731216

**Authors:** Susan D’Costa, Matthew J. Rich, Brian O. Diekman

## Abstract

Risk factors for the development of osteoarthritis (OA) include genetic background and focal cartilage injury. The search for disease-modifying OA therapies would benefit from a more comprehensive knowledge of the genetic variants that contribute to chondrocyte dysfunction and the barriers to cartilage regeneration. One goal of this study was to establish a system for producing engineered cartilage tissue from genetically-defined primary human chondrocytes through genome editing and single-cell expansion. This process was utilized to investigate the functional effect of bi-allelic knockout of the cell cycle inhibitor p21. The use of ribonucleoprotein (RNP) CRISPR/Cas9 complexes targeting two sites in the coding region of p21 resulted in a high frequency (16%) of colonies with homozygous p21 knockout. Chondrogenic pellet cultures from expanded chondrocytes with complete loss of p21 produced more glycosaminoglycans (GAG) and maintained a higher cell number. Single-cell derived colonies retained the potential for robust matrix production after expansion, allowing for analysis of colony variability from the same population of targeted cells. The effect of enhanced cartilage matrix production in p21 knockout chondrocytes persisted when matrix production from individual colonies was analyzed. Chondrocytes had lower levels of p21 protein with further expansion, and the difference in GAG production with p21 knockout was strongest at early passages. These results support previous findings that implicate p21 as a barrier to cartilage matrix production and regenerative capacity. Further, this work establishes the use of genome-edited human chondrocytes as a promising approach for engineered tissue models containing user-defined gene knockouts and other genetic variants for investigation of OA pathogenesis.

## Introduction

Osteoarthritis (OA) is characterized by joint pain and the progressive degradation of articular cartilage and surrounding tissues.^1, 2^ Primary human chondrocytes are a valuable resource for understanding the molecular pathways involved in cartilage dysfunction and OA, but the tools for manipulating chondrocytes have been mostly limited to partial knockdown, overexpression, and pharmacological inhibition. Genetics contribute approximately half of the risk for OA^3^ and there is a need to develop methods for the functional characterization of genetic variants that have been identified by genome-wide associated studies (GWAS) studies.^4-6^. The emergence of genome editing techniques provides an opportunity to knock out genes of interest and model genetic variants in chondrocytes.^7^ However, one limitation of genome editing is that the stochastic nature of editing outcomes necessitates the expansion of single-cell derived colonies in order to obtain genetically-defined cell populations. While this approach is compatible with immortalized cell lines and induced pluripotent stem cells (iPSCs), it presents challenges for primary cell types that would otherwise be ideally suited for the research question. One goal of this study was to establish a system by which primary human chondrocytes could be edited and expanded as individual colonies for subsequent cartilage tissue engineering.

The poor healing capacity of articular cartilage has spurred the development of numerous approaches to provide compensatory regeneration for focal cartilage defects, including marrow stimulation by microfracture^8^, autologous chondrocyte implantation^9^, and tissue engineering^10^. Understanding the blocks to endogenous regeneration will improve these treatment paradigms and may also lead to new strategies for the treatment and prevention of OA, as work with a set of recombinant inbred mice has connected the lack of regenerative capacity to OA susceptibility. Genetic backgrounds conducive to superior healing of ear holes have increased articular cartilage regeneration^11^ and are protected from OA.^12, 13^ One mediator of cartilage healing is the cell cycle inhibitor p21, as p21 knockout mice were able to recapitulate the ear closure regenerative phenotype of Murphy-Roths-Large (MRL) “superhealer” mice.^14^ Our previous work used short-hairpin RNA knockdown of p21 in iPSC-derived chondrocytes to demonstrate that reducing the level of p21 enhanced the synthesis of cartilage matrix.^15^ This finding is consistent with the relationship between p21 levels and differentiation potential of synovial mesenchymal stem cells,^16^ as well as the enhanced chondrogenesis with pharmacologic inhibition of p21 in this cell type.^17^. The current study generates engineered tissue from edited primary human chondrocytes in order to investigate the cartilage-forming potential of cells with and without the complete knockout of p21.

## Materials and Methods

### Isolation and culture of human articular chondrocytes

Normal human cartilage was obtained from ankle joints without OA from three male donors aged 49 (donor 1), 47 (donor 2), and 50 (donor 3) years old through the Gift of Hope Organ and Tissue Donor Network (Itasca, IL) via Rush Medical College (Chicago, IL). Primary human articular chondrocytes were isolated by sequential pronase and collagenase digestion as previously described^18^. Chondrocytes were plated at a density of 1.5 million cells per 6 cm^2^ dish (∼70,000 cells/cm^2^) in chondrocyte media consisting of Dulbecco’s Modified Eagle Medium/Nutrient Mixture F-12 HEPES (11330-032, Gibco) supplemented with 10% FBS (cat # 1500-500, VWR), 1x Penicillin-Streptomycin (15140122, Gibco), 2.5 µg/ml Amphotericin B (A2942, Sigma) and 4 µg/ml Gentamicin (1575-060, Gibco). Following recovery for 8-10 days, cells were trypsinized (T3924, Sigma), washed twice with PBS, and centrifuged at 100 g for 8 min in preparation for transfection.

### Preparation of genome editing components

Ribonucleoprotein (RNP) complex containing the Cas9 enzyme and sequence-targeting guide RNAs were prepared according to manufacturer’s recommendation (Integrated DNA Technologies). Briefly the tracrRNA (1072533, IDT) and crRNA (Human p21 guide AD: CCAAGCTCTACCTTCCCACG, Human p21 guide AC: TTCTGACGGACATCCCCAGC, Negative Control crRNA #1) were each resuspended in Tris-EDTA buffer to 200 μM stock concentration. Equimolar concentrations of crRNA and tracrRNA was combined and heated at 95 °C for 5 min and then cooled to produce the 100 μM crRNA:tracrRNA duplex. Separate RNP complex for each guide was prepared by combining the crRNA:tracrRNA duplex with 10 µg/µl Alt-R® Cas9 Nuclease (1081058, IDT) and PBS at a ratio of 1.2 : 1.7 : 2.1 at room temperature for 15 min.

### Transfection of primary human chondrocytes with RNP complex

RNP complex was delivered using the Amaxa Nucleofector™ system (Lonza). Following trypsinization and washing with PBS, approximately 200,000 cells were resuspended in a mixture containing 20 µl of P3 Primary Cell Nucleofector™ solution (V4XP-3032, Lonza), 5 µl of RNP complex for each guide (both guides AC and AD, or negative control), and 1 µl of 100 µM Alt-R® Cas9 Electroporation Enhancer (1075916, IDT). The mixture was gently pipetted up and down and transferred to the 16-well Nucleocuvette™ strip (V4XP-3032, Lonza) and transfected using program ER-100 on a 4D-Nucleofector™ Core unit (Lonza). Transfected cells were incubated at room temperature for 5-10 min and then transferred to 24-well plate containing prewarmed antibiotic free chondrocyte media supplemented with 20% FBS for recovery. An aliquot of the cells was placed in a 96-well to be used for confirmation of successful genome editing by PCR. Following confirmation of editing in the bulk cell population of the 96-well plate (2-4 days), the cells from the 24-well plate were passaged and seeded at low cell density (200 cells per 6 cm^2^ dish) for 8-12 days of expansion as single-cell derived colonies. Notably, colonies formed as seen in previous work^19^ despite the difference that the current approach did not perform fibronectin-based selection of a chondrocyte sub-population^19^. Individual colonies were picked with a pipette tip using a microscope (EVOS FL, ThermoFisher) placed in a laminar flow hood and split into 96-well and 24-well plates for analysis of genome editing and continued expansion, respectively.

### Screening single-cell derived colonies for editing outcomes by PCR and sequencing

Cells in 96 well plates were washed once with PBS before addition of QuickExtract™ DNA Extraction Solution (Lucigen). The volume was adjusted based on the estimated cell number, with a minimum of 35 µl and maximum of 100 µl for a nearly confluent well. After a 15 minute incubation at 37 °C, cells were pipetted up and down, transferred to tubes (490003-722, VWR), and vortexed for 1 minute. The cell suspension was then heated at 65 °C for 6 minutes, vortexed briefly, and placed at 98 °C for 2 minutes. Extracted DNA solution was kept at -20 °C until PCR. PCR amplification was performed by adding 5 µl template DNA, 1 µM forward (TTAGCTTGCCCTTCAGTTGC) and reverse (5’-GGTCTTTGCTGCCTACTTGC-3’) primers, and EconoTaq PLUS GREEN 2X Master Mix (Lucigen) in a 25 µl reaction. PCR conditions included an initial denaturation at 94 °C for 2 minutes, 35 cycles of denaturation at 94 °C for 30 seconds, annealing at 58 °C for 30 seconds, and extension at 72 °C for 50 seconds, followed by a final extension at 72 °C for 10 minutes. For sequencing of PCR products, the DNA from the PCR reaction was purified using a clean-up kit (T1030, New England Biolabs) and sequenced using SimpleSeq™ tubes (Eurofins Genomics) and the following primer: ACCAGCTGGAAGGAGTGAGA. Chromatograms were visualized using BioEdit software.

### Cell expansion in monolayer

Selected colonies with confirmed edits (or negative controls) from the 24-well plate were trypsinized and plated as passage 1 cells at a density of ∼3,000 cells/cm^2^ in media supplemented with 1 ng/ml TGF-β1 (PHG9214, Life technologies) and 5 ng/ml bFGF (PHG0264, Life technologies). Pellets were made at the end of passage 3 (donor 1), the end of passage 1 (donor 2), or at the end of passage 1 through passage 4 (donor 3). The fold increase and number of population doublings were calculated based on counts obtained using a Countess™ II Automated Cell Counter.

### Pellet culture for evaluation of chondrogenic potential

Pellets were made from pooled colonies after expansion (donors 1 and 3) or from individual single-cell derived colonies (donor 2). Briefly, 250,000 cells were placed in 15 ml tubes and washed with serum free Dulbecco’s Modified Eagle Medium-high glucose (DMEM-HG, 11995-065, Sigma). DMEM was replaced with differentiation media consisting of DMEM-HG supplemented with 1% ITS+ Premix (354352, Corning), 1x Penicillin-Streptomycin (Gibco), 100 nM dexamethasone (D4902, Sigma), 50 µg/ml proline (P5607 Sigma), 50 µg/ml L-ascorbic acid 2 phosphate (A8960, Sigma), 1% FBS, and 10 ng/ml TGF β1. Cells were centrifuged at 200 g for 10 min and caps were loosened to permit gas exchange for culture in a standard incubator. Media was changed every two to three days and pellets were harvested on day 28. Pellets were blotted dry and weighed before subsequent analysis.

### Analysis of total DNA and glycosaminoglycan (GAG) content

Pellets were digested with a solution consisting of 125 μg/mL papain, 0.1 M sodium phosphate, 5 mM EDTA and 5 mM cysteine hydrochloride at pH 6.5. Digestion was performed for 24 hours at 65 °C in an oven or with agitation in a ThermoMixer C (Eppendorf) at 600 rpm. The QuantiFluor® dsDNA System was used to measure dsDNA content according to manufacturer’s instructions. GAG content of the pellet was evaluated using the DMMB assay as previously described^20^ with 1,9 dimethylmethylene blue dye at pH 3 and a chondroitin-4-sulfate standard curve.

### Histology and Immunohistochemistry

Pellets were fixed in 4% paraformaldehyde for 24 hours at 4 °C and processed for paraffin embedding. For histology, slides were stained with Fast Green (0.02% in dH20, 25 minutes, Sigma 7252) and Safranin-O (1.5% in dH20, 30 minutes, Sigma S8884). For immunohistochemistry, antigen retrieval was performed using pepsin (Digest-All™ 3, ThermoFisher 003009) for 5 minutes at room temperature. Samples were incubated overnight at 4 °C using a primary antibody targeting type II collagen (II-II6B3, Developmental Studies Hybridoma Bank, no dilution), followed by an anti-mouse secondary antibody (715-065-151, Jackson ImmunoResearch, 1:1000) and visualization using VECTASTAIN® Elite ABC-HRP (PK-6100, Vector Laboratories) and DAB Peroxidase (SK-4100, Vector Laboratories). For both histology and IHC, slides were counterstained with hematoxylin and mounted with Permount™ for imaging.

### Protein isolation and western blotting

Protein isolation and western blotting were performed as described^21^. Briefly, TRIzol™ (ThermoFisher) was used on both monolayer cells and pellets. For monolayer, TRIzol™ was added to cells and vortexed for 1 minute. Pellets were placed in bead beater tubes (10158-610, VWR) with TRIzol™ and homogenized at 6500 rpm × 30 seconds for 3 cycles (Precellys^®^ 24, Bertin Corp). Protein concentration was determined using the Micro BCA Protein assay kit (Thermo Fisher) and either 12 µg (monolayer cells) or 8 µg (pellets) was used for SDS-PAGE. The membrane was cut after blocking and the lower half was incubated in p21 antibody (2947 Cell signaling Technology 1:2000) while the upper half was incubated in anti-G3PDH Human Polyclonal Antibody (GAPDH) (2275PC, Trevigen 1:2000), both overnight at 4 °C. Following incubation in secondary antibody solution (Cell signaling technology, 1:2000) for one hour, the membranes were washed and placed in the Radiance Plus chemiluminescent substrate and imaged with the Azure c600 gel imaging system (Azure Biosystems).

### Statistical analysis

Statistical analysis and plotting was performed using Prism 7 (GraphPad, La Jolla, CA, USA). Data are plotted as individual points and error bars indicate mean ± standard error of the mean (SEM). Normality was assessed by Shapiro-Wilk test and data were analyzed by unpaired t-test or Two-way ANOVA with Tukey’s post-hoc analysis.

## Results

### Genome editing in primary human chondrocytes

We developed an efficient system for generating engineered tissue from primary human chondrocytes with the homozygous deletion of a target gene. This process is summarized in Figure 1 and involves delivering RNP complexes for CRISPR/Cas9 based editing, screening single-cell derived colonies, and then generating chondrogenic pellet cultures after further expansion. The genomic DNA of p21 was targeted with guide RNAs that generate double-stranded breaks at 15 base pairs and 237 base pairs into the coding region (Fig. 2A). Thus, cutting at both locations would generate an approximately 222 base pair deletion. The bulk population of edited cells was assessed by PCR amplification of the targeted region, which showed that including the p21 guide RNAs generated a novel amplicon that is ∼222 base pairs smaller than the parent product (Fig. 2B, left). Repeating this analysis on single-cell derived colonies allows for the discrimination of the editing outcome in each colony: fully intact (parent product only), heterozygous targeted (both parent and smaller product), and homozygous targeted (smaller product only) (Fig. 2B, right). The efficiency of editing was estimated by analyzing the percentage of screened colonies with each editing outcome. Analysis of 80 colonies from donors 2 and 3 showed that 53 (66%) colonies had a heterozygous deletion and 13 (16%) had a homozygous deletion (Fig. 2C). To gain further insight into the exact editing outcomes, we performed Sanger sequencing on a subset of the PCR products (Fig. 2D). The homozygous colonies confirmed a single read with the expected deletion, with the example shown having the same 221 base pair deletion on both alleles (Fig. 2D). Notably, even the colonies scored as “wild type” by PCR due to the lack of a large deletion showed multiple reads beginning at the expected guide cut site, which is consistent with the error-prone non-homologous end joining (NHEJ) repair pathway (Fig. 2D).

**Figure 1.**
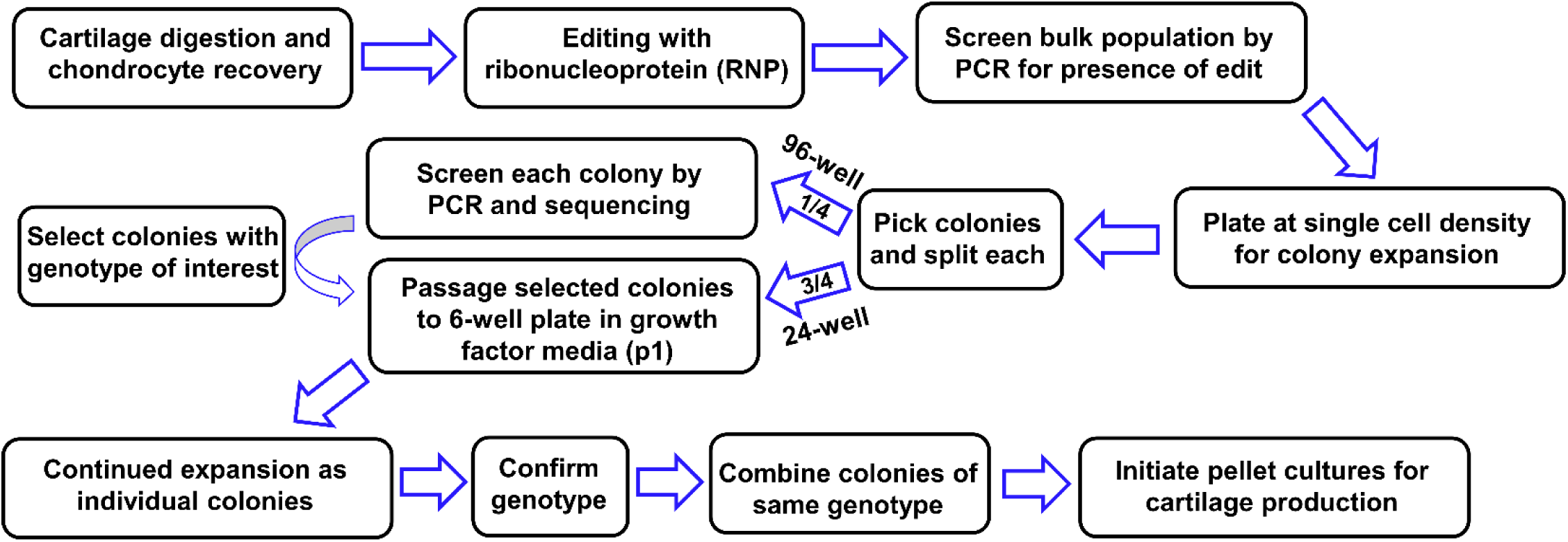
Workflow of system for engineering cartilage tissue from genome-edited primary human chondrocytes.

**Figure 2.**
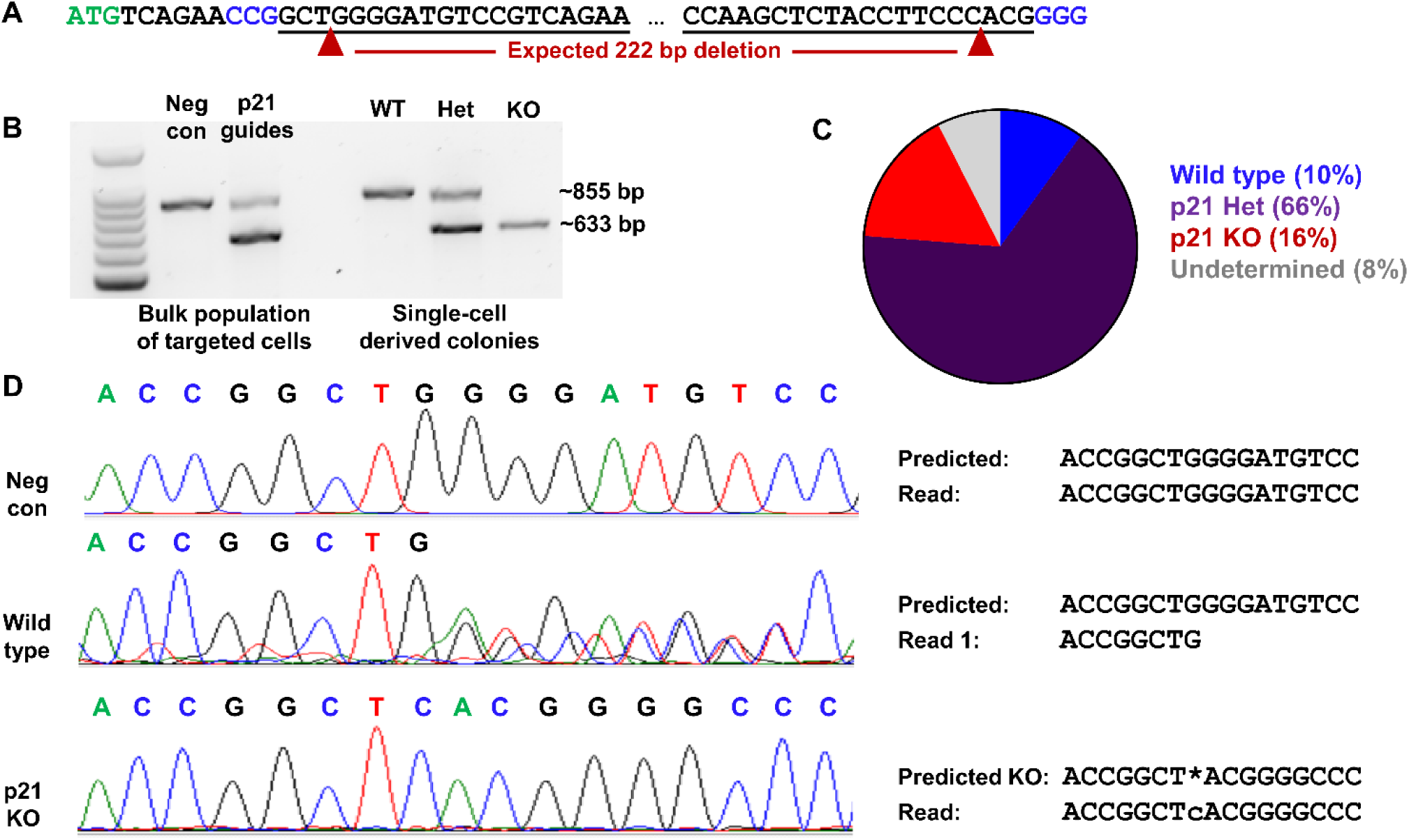
Screening and efficiency of homozygous p21 knockout. A) Sequence of p21 targeted by editing, with start of coding region in green, guide RNA target sites underlined, PAM sites in blue, expected cut sites denoted by red arrows. B) Representative gel showing the emergence of the ∼222 base pair shorter product with the inclusion of guide RNAs targeting p21. DNA from bulk population of targeted cells (left) or individual colonies (right). C) 80 colonies from donors 2 and 3 were screened for editing outcomes by PCR. D) Sanger sequencing of representative colonies with predicted and actual reads noted.

### Effect of homozygous p21 knockout on pellet chondrogenesis (donor 1)

Individual colonies that had been targeted by the negative control guide RNA (p21 intact) and those screened and sequenced as p21 knockout were expanded in either 10% FBS chondrocyte medium or with the addition of 1 ng/ml TGF-β1 and 5 ng/ml bFGF. The growth factors resulted in a larger fold increase during passage 1 regardless of genotype (Fig. S1A). As engineered tissue production requires a substantial number of cells and previous studies have shown that culture with TGF-β1 and bFGF is conducive to subsequent chondrogenesis ^19, 22^, further studies were completed using the growth factors for expansion. Under these conditions, all colonies from p21 intact and p21 knockout groups were capable of more than 22 population doublings from the time of single-cell plating through the end of the third passage (Fig. S1B), which is equivalent to over 4 million cells per starting colony. To determine the effect of p21 knockout, we first combined the expanded colonies of each genotype and confirmed the complete loss of measurable p21 protein by western blot (Fig. 3A). Chondrogenic pellet cultures with 250,000 cells each were established from both genotypes and those made from p21 knockout chondrocytes had a higher weight at the end of the 28-day culture period (Fig. 3B). Digestion of the pellets for analysis of biochemical content showed that p21 knockout pellets also had significantly higher GAG content (Fig. 3C). Part of this increased matrix production is likely due to the higher number of cells in pellets from p21 knockout chondrocytes as assessed by double-stranded DNA content (Fig. 3D). Paraffin sections were analyzed to show the presence of enhanced glycosaminoglycans (Fig. 3E) and type II collagen (Fig. 3F) in the p21 knockout pellets as compared to controls.

**Figure 3.**
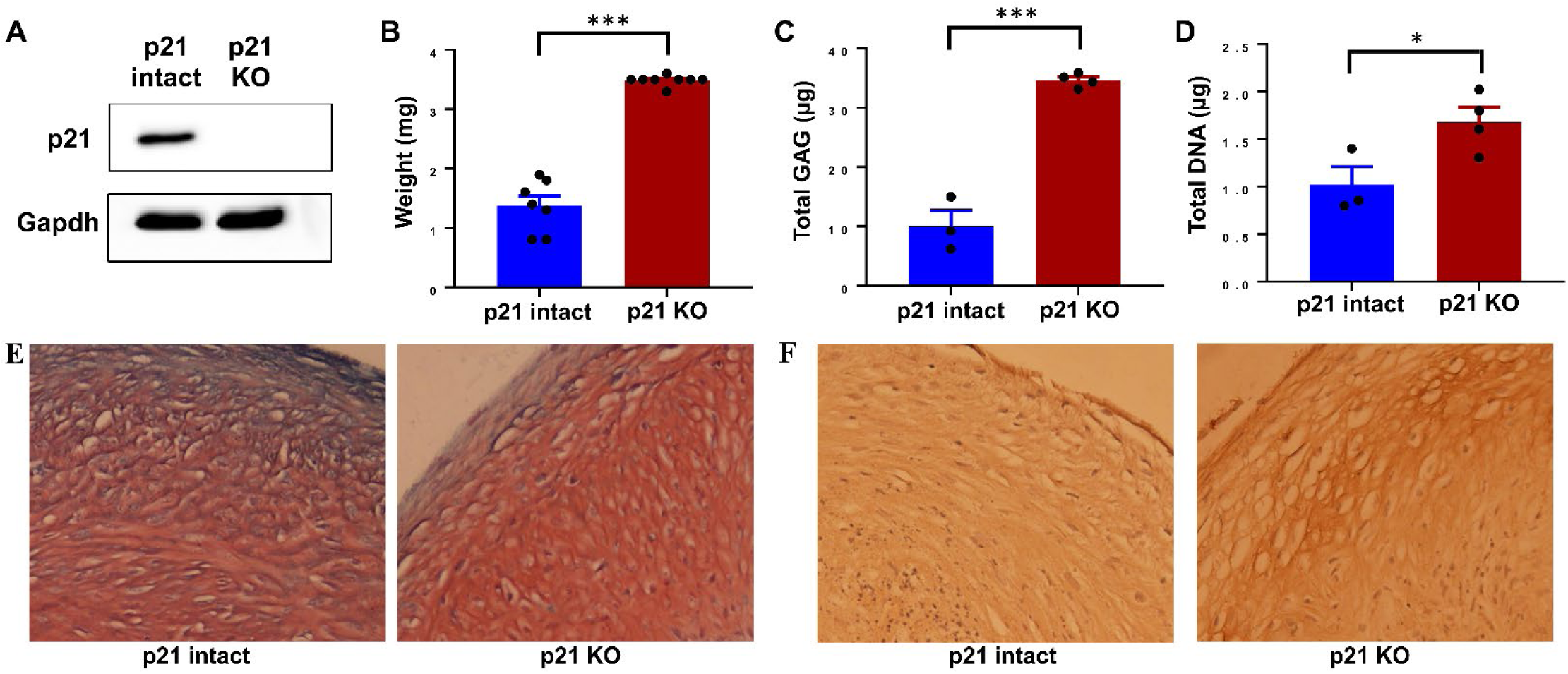
Cartilage matrix production with p21 knockout (donor 1). A) Western blot analysis of combined colonies genotyped as p21 intact or p21 knockout (KO). B) Pellet weight after 28 days of culture. C) Glycosaminoglycans (GAG) per pellet. D) DNA per pellet as a surrogate for cell number. E) Histological sections (red: Safranin-O, green: Fast Green, purple: hematoxylin) imaged at 20x. F) Immunohistochemistry for type II collagen (brown) with hematoxylin counter stain (purple), imaged at 20x. * indicates p<0.05, ** indicates p<0.01, *** indicates p<0.001.

### Differences in chondrogenic potential by colony (donor 2)

Combining the cells of expanded colonies before pellet culture eliminates the potential to investigate the variability in how a particular edit (in this case p21 knockout) affects chondrogenesis. Thus, edited chondrocytes from donor 2 were expanded through end of the first passage and made into pellet cultures from 10 p21 intact and 5 p21 knockout colonies. Biochemical data from independent pellets for each colony were typically clustered for both GAG (Fig. 4A) and DNA (Fig. 4B), with larger variation between different colonies of the same genotype. For clarity, data were also averaged to a single value per colony for both GAG (Fig. 4C) and DNA (Fig. 4D). The variability between colonies from the same bulk population of cells is consistent with previous work analyzing the chondrogenic potential of single-cell derived chondrocyte colonies.^22, 23^ However, the effect of p21 knockout persisted through the colony variability, as both DNA content and GAG content were increased in the set of colonies with homozygous loss of p21 as compared to control.

**Figure 4.**
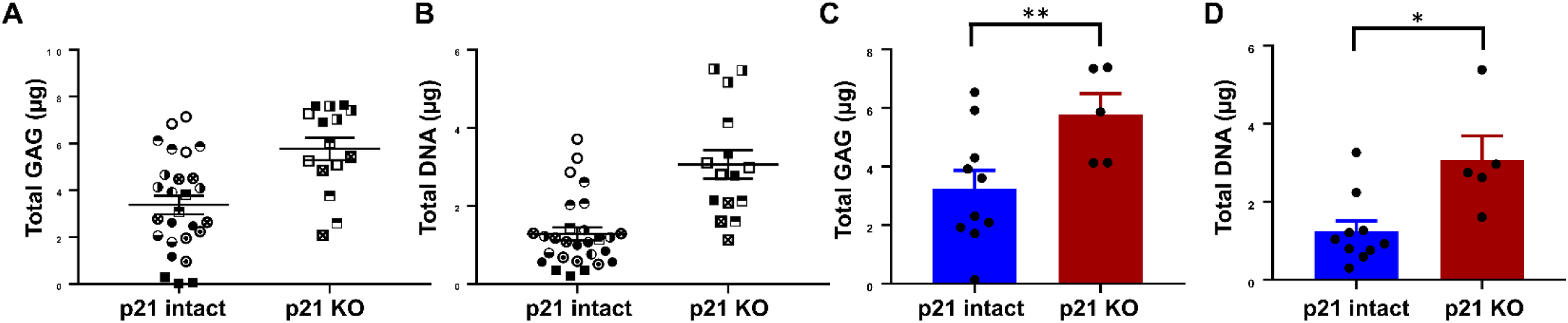
Pellets from individually expanded colonies (donor 2). For panels A and B, matched symbols indicate replicate pellets from the same expanded colony. For panels C and D, the mean value for each colony is plotted and used as an independent value for statistical analysis. A, C) Glycosaminoglycans (GAG) per pellet. B, D) DNA per pellet as a surrogate for cell number. * indicates p<0.05, ** indicates p<0.01.

### The effect of p21 knockout on chondrogenesis of expanded chondrocytes (donor 3)

Reducing the level of p21 through hypoxia, growth factors, or pharmacological inhibition has been shown to maintain proliferative and differentiation potential during the expansion of mesenchymal stem/stromal cells (MSCs).^17, 24-26^ To determine the way in which p21 knockout and continued expansion affect chondrogenesis, we established pellet cultures at the end of passages 1 through 4. As expected given the genetic nature of the edit, the complete knockout of p21 was confirmed to be stable both during expansion and throughout pellet culture (Fig.5A). Protein levels in the p21 intact samples were quantified and the production of p21 was similar at the end of pellet culture as compared to monolayer cells at that passage (Fig. 5B). The level of p21 protein decreased with extensive passaging (Fig. 5B), perhaps due to repression by the presence of bFGF or due to preferential expansion of p21-low chondrocytes within the population. Analysis of the pellets at the end of each passage demonstrated a significant effect of both passage number and genotype by Two-way ANOVA for pellet weight (Fig. 5C), GAG content (Fig. 5D), and DNA content (Fig. 5E). Comparisons between genotypes at each passage indicated that the p21 knockout had the strongest effect on pellets made at passage 1, with a significant effect on pellet weight, DNA content, and GAG content. While in this donor, the chondrogenic potential of cells at later passages was not affected by the knockout of p21 (Figures 5C-E).

**Figure 5.**
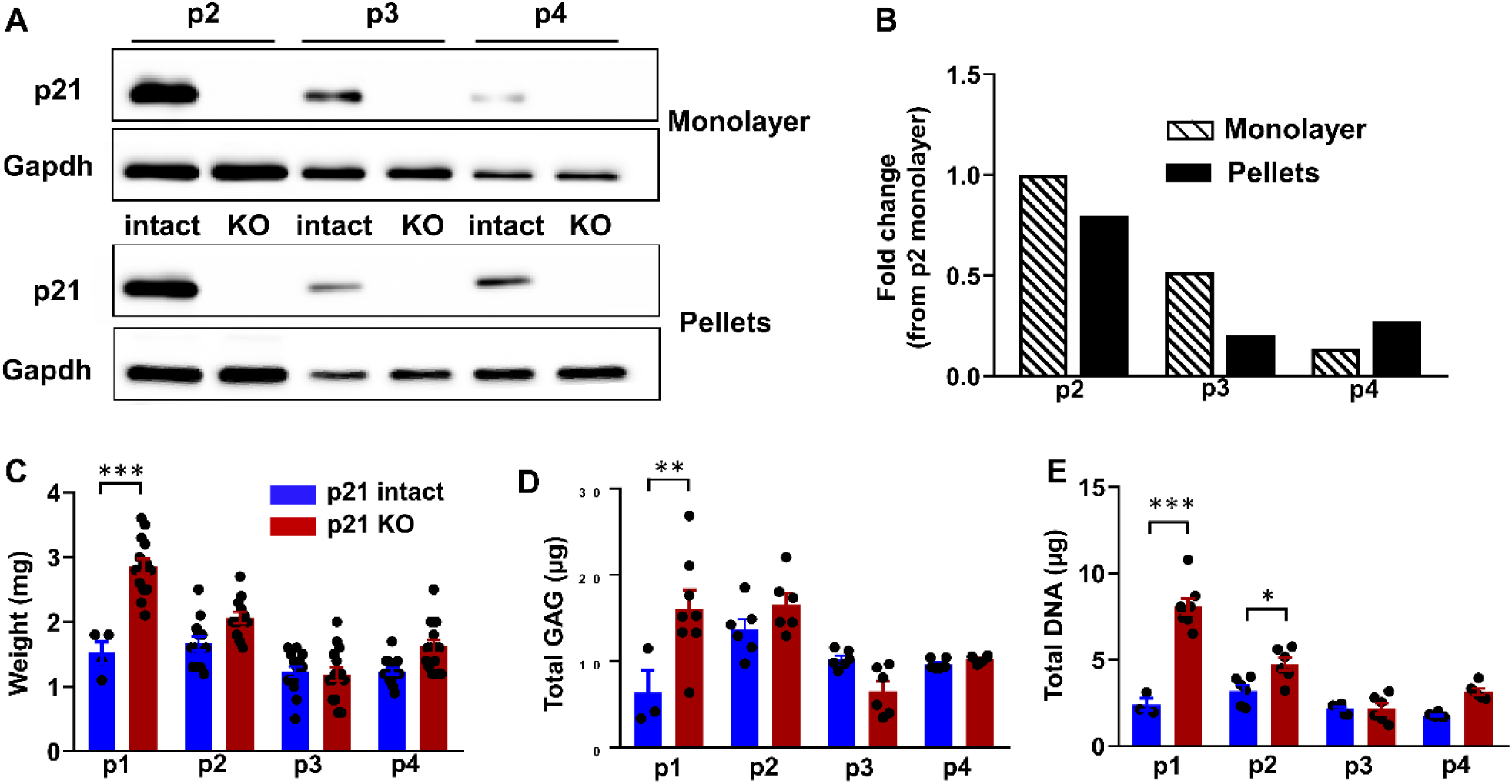
Effect of continued expansion in monolayer (donor 3). A) Western blot analysis of cells in monolayer culture (top) and pellets after 28 days of culture (bottom) at passages 2, 3, and 4 with or without p21 knockout (KO). B) Quantification of western blot pixel intensity, normalized to Gapdh and then to passage 2 monolayer cells. C) Pellet weight after 28 days of culture. D) Glycosaminoglycans (GAG) per pellet. E) DNA per pellet as a surrogate for cell number. * indicates p<0.05, ** indicates p<0.01, *** indicates p<0.001.

## Discussion

The ability to establish genetically-defined populations of primary human articular chondrocytes has profound implications for the study of cartilage biology and OA.^7^ However, a major barrier to genome editing in chondrocytes is the challenge of expanding edited colonies without losing the potential to synthesize robust cartilage matrix. This study establishes an efficient approach for engineering cartilage tissue from primary human chondrocytes and validates the method by targeting the cell cycle inhibitor p21. Consistent with the hypothesized role of p21 in restraining regenerative capacity, cartilage tissue generated from chondrocytes with complete p21 knockout had more cells and a higher GAG content.

This work is complementary to other genome editing approaches that have been used to study cartilage biology. Yang et al expressed Cas9 in a rat chondrosarcoma cell line to allow for the knockout of genes through guide RNA transfection.^27^ The authors then generated stable cell lines with complete knockout of aggrecan to demonstrate the effect of this critical cartilage component on cell morphology and chondrosarcoma development.^27^ Seidl et al targeted a bulk population of primary human chondrocytes with the RNP version of CRISPR/Cas9 to initiate a reduction in functional MMP-13.^28^ While single-cell derived colonies were not selected in this study, generating NHEJ in the coding region of MMP-13 was sufficient to reduce the level of MMP-13 in the bulk cell population. Because MMP-13 is a secreted factor that compromises function at the tissue level, the reduced MMP-13 resulted in a higher quality engineered tissue.^28^ Another genome editing strategy is to modify pluripotent cells before subsequent differentiation to the chondrogenic phenotype. While using iPSC-derived cartilage for modeling OA is attractive in terms of storing edited progenitors and the opportunity for tissue scale-up,^29^ genome modification occurs during the pluripotent stage and thus the effect of the change may be enacted during the complex differentiation towards the chondrocyte phenotype.^30-33^ Each of these previous approaches (generating stably modified cell lines, bulk targeting of chondrocytes, iPSC editing and differentiation) has merit for particular applications and we propose that the current paradigm will be particularly useful in experiments that require genetically-defined populations of primary articular chondrocytes.

We selected p21 as a first target for this approach due to the intriguing body of literature suggesting that p21 is a potent repressor of tissue regeneration. The Murphy-Roths-Large (MRL) “superhealer” mouse strain was originally identified by the scarless closure of ear holes.^34^ MRL mice also showed improved regeneration in many other tissue contexts, including full-thickness articular cartilage defects.^35^ Following the lead of altered cell cycle distribution in cells from MRL mice, Bedelbaeva et al showed that the regenerative capacity was linked to lower inducible levels of p21 and that p21 knockout mice showed the ear hole closure phenotype^14^. Subsequent lineage tracing studies in p21 knockout mice demonstrated that regenerated cartilage is produced by previously existing chondrocytes that re-enter the cell cycle and produce new tissue.^36^ Interestingly, p21 knockout mice are more susceptible to an instability model of post-traumatic OA, illustrating the complex nature of disease pathogenesis and raising the possibility that changes to other aspects of OA may counter the potentially beneficial effects of enhanced cartilage regenerative potential.^37, 38^

Given the complexity of in vivo regenerative capacity, in vitro models can be useful as a way to focus on tissue-intrinsic changes with loss of p21. We had previously shown that the stable (but incomplete) shRNA knockdown of p21 in murine iPSC-derived chondrocytes increased the chondrogenic potential of this cell type.^15^ The current use of permanent (and complete) knockout in primary human chondrocytes gave similar results in terms of pellet cultures with increased weight, cell number, and GAGs. However, the interaction of p21 loss and cell expansion on chondrogenic potential was different with the two cell types. In the previous study, p21 knockdown cells generated similar quality engineered cartilage pellets at the end of 2 or 5 passages, which was different than control cells and therefore extended the expansion window before the loss of chondrogenic potential.^15^ In the current study, p21 knockout had the largest effect on cells at early passages and p21 knockout cells at the end of passage 4 were similar to controls.

One limitation of the single-cell colony approach is that heterogeneity of the outcome between individual colonies of the same genotype may make it challenging to detect the effect of the genome editing. The potential variability is illustrated by Fellows et al in their description of the number of population doublings before senescence in chondrocyte colonies from normal and OA patients.^19^ Colony variability in expanded chondrocytes can also lead to differing capabilities for cartilage matrix production.^22, 23^ To explore the effect of colony variability in our system, we made pellet cultures from individually expanded colonies of each genotype (donor 2, figure 4). While there was differing chondrogenic capacity in these colonies, the genotype effect of p21 knockout was still present as evidenced by a significant increase in GAG content and DNA. This reinforced the data from donors 1 and 3 in which pellets were made from the combined cells of numerous colonies of each genotype at the end of expansion. These findings suggest that other applications of this technology will need to consider colony variability during experimental design, especially when the genetic modification is expected to generate a relatively subtle effect on matrix production.

A major contribution of this work is to establish a framework for genome editing in primary human chondrocytes. One particularly exciting use of this technology is for the functional follow-up of OA risk variants. Follow-up from GWAS studies has required techniques such as allelic expression imbalance and reporter assays to determine the function of the specific genetic changes.^39, 40^ The current approach extends the possibilities of functional validation by generating isogenic populations of primary chondrocytes - where the only difference from control cells is the user-defined change. These sophisticated models will be critical for assessing the increasing number of genetic variants that have been identified as risk factors for OA.

## Acknowledgments

The authors would like to thank the UNC Animal Histopathology core. This work was conducted while Brian Diekman was an Arthritis and Aging Research Grant recipient from the Arthritis National Research Foundation (ANRF) and the American Federation for Aging Research as well as second-year funding from ANRF.

## Figures and Figure Legends

**Figure S1:**
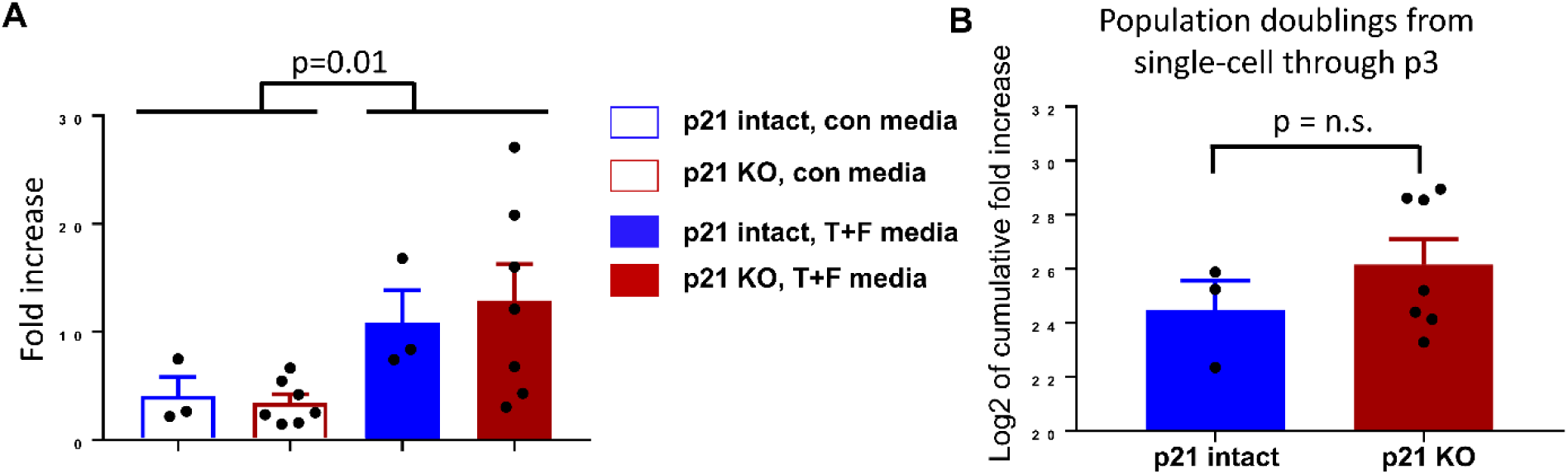
Expansion of single-cell derived colonies (donor 1). A) Colonies with p21 intact or p21 knockout (KO) were split to control conditions (10% serum, no growth factors) or media containing 1 ng/ml TGF-β1 and 5 ng/ml bFGF (T+F media). The fold increase over first passage (4 days) was calculated by cell counting. B) The cumulative number of population doublings from the time of plating at single-cell density to the end of passage 3 was calculated for colonies expanded in T+F media.

